# Determination of sex of sacrum in adult *Macaca mulatta*^1^

**DOI:** 10.1101/428748

**Authors:** Yan-Mei Wang, Xiao-Fan Han, Xiao-Jin Zhao

## Abstract

**Background:** Sacrum being a part of pelvis is an important bone for identification of sex in both living primates and fossil ones.

**Aim:** Aim of this work was to examine the sex differences of sacral parameters in rhesus macaques and to compare with those of the other primates.

**Materials and Methods:** Fifty-six adult scara of macaques (17 males and 39 females) have been investigated. Measurement of various parameters was done using sliding vernier calliper; and statistical analysis was done using SPSS 23.0 package.

**Results:** The present study showed that the cranial breadth of the sacrum, the sacral length, transverse diameter and sagittal diameter of the cranial articular surface, and two indices of relative sacral breadth were highly significant for sex determination in *Macaca mulatta*. Comparison of the present data with other studies suggest that sex determination of sacrum can be very different in various types of primates.

**Conclusion:** The results suggest that these measures may be functionally integrated in response to locomotion, obstetric adequacy and cephalopelvic proportions in primates. Sacral index is more reliable and should be applied for sex determination of sacrum in various anatomical and anthropological investigations.

---

Sex determination from bone plays a useful role in anatomy and anthropology personnel. Sacrum is the most commonly used bone for sex identification, as has been amply demonstrated in human^[1–3]^. Do these differences represent a human peculiarity, or are they comparable in character and degree to sex differences in the sacrums of nonhuman primates?

In a special publication on sex differences in pelves of monkeys and apes, Tague et al. concluded that practically all the pelvic characters which tend to separate the sexes in man show more or less marked sex differentiation in the nonhuman primates^[4,5]^.

Zhao et al. reported that the secondary sex differences were very pronounced in the pelvis of adult *Macaca mulatta* from a population in the Taihang Mountains in northern China^[6–8]^. Despite long-standing interest in sexual dimorphism of the human sacrum, comparatively few studies of sacral sexual dimorphism in nonhuman primates have been conducted.

Present study was carried out to assess the efficacy of the variables of sacrum in the *Macaca mulatta* to determine sex.

## Materials and Methods

Basic data for this study has been gained on specimens of adult Macaca mulatta, which were collected from the Taihang Mountains in Henan province by Dr. Zhao Xiaojin and now housed at the College of Life Science, Henan Normal University. Animals died from natural factors or injury between the ages of 5 and 25 years were selected to use. Sex was determined before the animals were skeletonized. Materials used in present study were 56 sacra (17 males & 39 females) of known sex. All the bones used were free of deformities or pathological changes in the study. All measurements were taken by one of the authors (ZXJ) and were recorded with a caliper to an accuracy of 0.01mm.

Measurements taken on each sacrum were illustrated in Fig. 1.

**Fig. 1.**
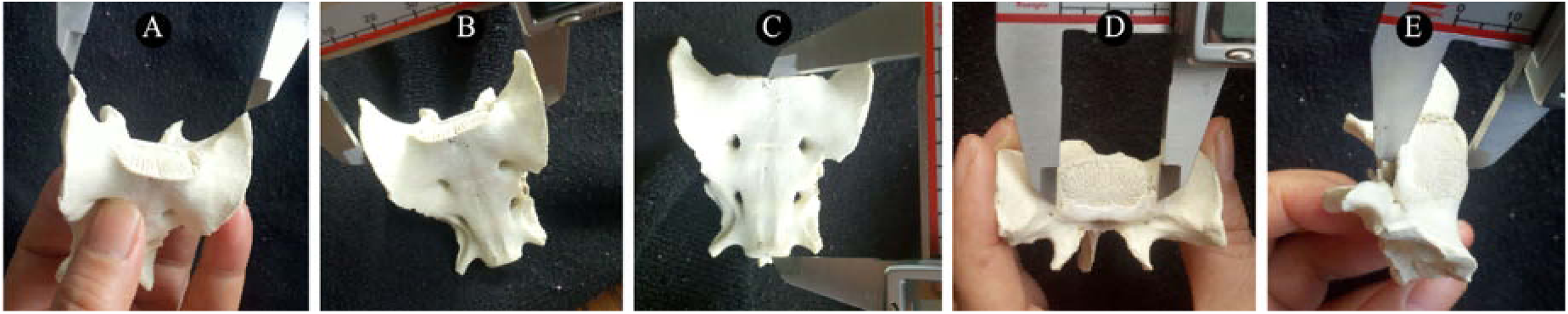
Dimensions of the sacrum measured in this study. A. Cranial breadth of the sacrum; B. Sacral breadth; C. Sacral length; D. Transverse diameter of the cranial articular surface; E. Sagittal diameter of the cranial articular surface.

The data presented were then subjected to statistical analysis using the SPSS for Windows (version 22.0).

Jit and Singh proposed that demarking points of various parameters were used for sex identification of sacrum^[8]^. They suggested for identification of male sacrum, the demarking point of a particular measurement was more than 3 SD. of mean value for female, and, for identification of female sacrum, the demarking point of same measurement was less than 3 SD. of mean value for male.

## Results

Summary statistics for each measurement were presented in Table 1.

**Tab. 1.**
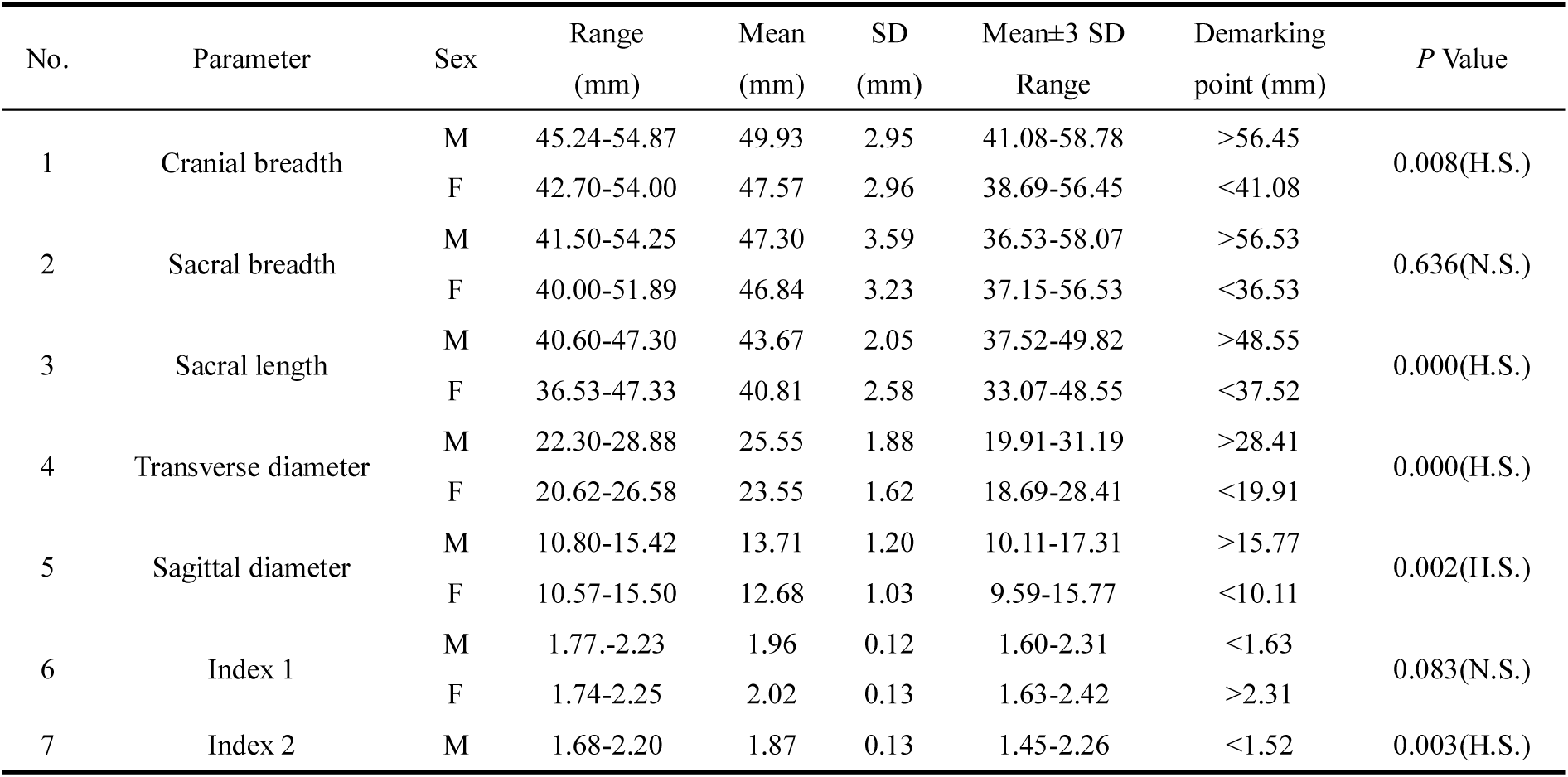
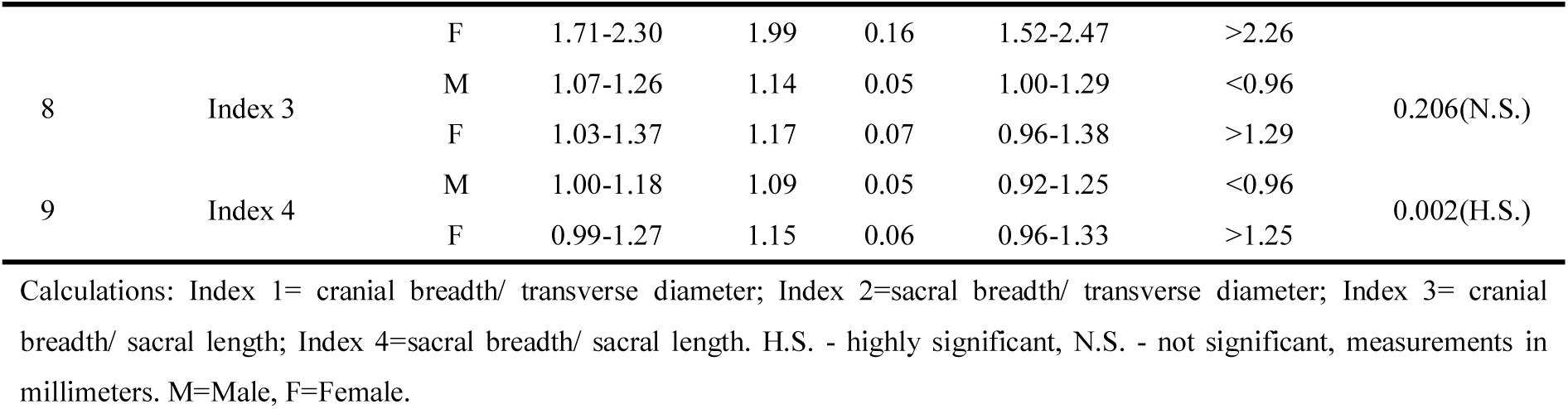
Showing various parameters of sacrum and their statistical analysis

In the present study, the parameters such as cranial breadth of sacrum, sacral length, transverse diameter of the cranial articular surface and sagittal diameter of the cranial articular surface were higher in males. And the values for parameters such as index 2 and index 4 of sacrum were higher in females.

For example, the mean value of sacral index 2 was 1.87 in males and 1.99 in females. Correspondingly, the demarking points in males was <1.52 and >2.26 in females using this parameter alone with high level of significance (*P*=0.003). The results were compared with the available literature in Table 2.

**Tab. 2.** Compared with previous studies for sexual dimorphism in various parameters of sacrum^a^

| Species | Category<sup>b</sup> | Sacral breadth |  |  | Transverse diameter |  |  | Sacral breadth/ transverse diameter |  |  |
| --- | --- | --- | --- | --- | --- | --- | --- | --- | --- | --- |
|  |  | M | F | P | M | Fe | P | M | F | P |
| *H. sapiens* | A | 107.13 | 108.68 | 0.335 | 52.84 | 47.59 | <0.001 | 2.02 | 2.27 | <0.001 |
| *H. lar* | A | 35.36 | 38.40 | <0.001 | 19.86 | 19.50 | 0.238 | 1.78 | 1.97 | <0.001 |
| *P. paniscus* | B | 60.87 | 61.45 | 0.823 | 31.45 | 31.69 | 0.849 | 1.96 | 1.93 | 0.991 |
| *P. troglodytes* | B | 63.45 | 64.75 | 0.594 | 35.79 | 34.5 | 0.377 | 1.93 | 1.90 | 0.221 |
| *N. larvatus* | C | 48.89 | 41.92 | <0.001 | 28.91 | 23.19 | <0.001 | 1.70 | 1.81 | 0.019 |
| *M. mulatta* | C | 47.30 | 46.84 | 0.636 | 25.55 | 23.55 | <0.001 | 1.87 | 1.99 | 0.003 |
| *G. gorilla* | D | 66.63 | 78.09 | <0.001 | 50.98 | 41.85 | <0.001 | 1.87 | 2.07 | <0.001 |
| *P. pygmaeus* | D | 78.12 | 74.14 | 0.297 | 39.04 | 35.57 | 0.150 | 2.02 | 2.03 | 0.412 |

<sup>a</sup> The data and the categories in the table were introduced by Moffet with the exception of *Macaca mulatta* in present study.
<sup>b</sup> The sampled taxa were divided into four categories based on distinct combinations of body size dimorphism and cephalopelvic proportions.
Categories defined as: (A) minimal body size dimorphism and large cephalopelvic proportions; (B) minimal body size dimorphism and small cephalopelvic proportions; (C) large body size dimorphism and large cephalopelvic proportions; and (D) large body size dimorphism and small cephalopelvic proportions.
M=Male, F=Female, P=P value

## Discussion

This paper is the first report describing the sexual dimorphism of the sacrum in *Macaca mulatta* from a population in northern China.

Schultz related pelvic sexual dimorphism among monkeys to obstetric factors, but argued body size dimorphism to be the primary influence on pelvic dimorphism among the other apes, including gibbons, which have large cephalopelvic proportions. His study presented the possibility that pelvic dimorphism in primates was not just related to obstetric, allometric or phylogenetic explanation, but rather a multifaceted relationship among these variables ^[10]^.

Compared to other primates in this study, Moffett et al. have found sexual differences in relative sacral breadth among catarrhine primates encompassing *Homo sapiens, Hylobates lar, Gorilla gorilla*, and *Nasalis larvatus* (Tab. 2).

For *Macaca mulatta*, the sex differences in the relative sacral breadth are clearly more marked, though much less pronounced than in *Homo sapiens, Hylobates lar and Gorilla gorilla*. The index of the macaque lies between the indices of some hominoid taxa such as *Homo sapiens*, and *Hylobates lar* on one side and of the *Nasalis larvatus* on the other side. *Macaca mulatta* in present study is characterized by both large body size dimorphism and large cephalopelvic proportions, the combination which does not exist among living hominoids, as has been found in *N. larvatus*^[5]^.

No unambiguous experimental manipulation yet exists to link sexual dimorphism in relative sacral breadth to body size dimorphism or cephalopelvic proportions, but a strong inferential case can be built from existing evidence.

Firstly, as in most primates, the sacrum of *Macaca mulatta*, which influences the shape and dimension of the birth canal, is interpreted within obstetric contexts. Although the obstetrical difficulty of *Macaca mulatta* does not wholly explain this relationship, a reasonable inference is that females with small pelvis experience more obstetric difficulties than females with large pelvis^[11]^.

Secondly, another factor affecting the sexual dimorphism of sacrum is differences in locomotor habit among different primates. Leutenegger concluded that many of the dimensions, such as the distance between the sacroiliac joint and the hip joint, influence both the dimension of the birth canal and locomotor characteristics ^[11]^. Thus the average size of the pelvic outlet and the locomotor constraints may vary in different species. Females of a species in which locomotor adaptations favored a relatively smaller outlet than males might show greater enlargement of these dimensions ^[12]^.

Thirdly, most previous studies had proposed that the presence of the birth canal influenced the female primate pelvis structure, and other studies have explained that pelvic dimorphism generated by allometric body size between the sexes. Karen et al. concluded that sexual dimorphism of body size was not an exclusive determinant but an important factor influencing pelvic dimorphism on the basis of their coefficients of allometry ^[13]^. The coefficients of allometry were similar in *Colobus, Presbytis*, and *Cercopithecus;* in these taxa mean ischial length was greater in male skeletons but pubic length showed either no significant increase in the larger males or mean pubic length was greater in the smaller female skeletons ^[14]^.

Finally, Moffet et al. argued that species with large cephalopelvic proportions had higher fertility, for the females with wide sacral alae could enlarge the pelvic inlet during the delivery^[5]^. However, gorillas, with small cephalopelvic proportions, were unlikely to have relative wide sacrum. This may interpret that sacral dimorphism in gorillas is attributable to selection for relatively narrow sacra in males rather than relatively broad sacra in females.

Overall, the results in keeping with previous studies suggesting that these measures may be functionally integrated in response to locomotion, obstetric adequacy and cephalopelvic proportions in primates ^[8,15]^.

The present study shows that the cranial breadth and the sacral length of the sacrum, transverse diameter and sagittal diameter of the cranial articular surface, and two indices of relative sacral breadth are highly significant for sex determination of sacrum in *Macaca mulatta*.

## Footnotes

^1^[**Foundation item**] A grant from the Science and Technique Foundation of Henan Province (172102310721) [**Biography**] Yan-Mei Wang (1977—), female (Han), Lu yi Henan province, Lecturer. Tel: 13782525279

